# Comparison of video-based and sensor-based head impact exposure

**DOI:** 10.1101/235432

**Authors:** Calvin Kuo, Lyndia Wu, Jesus Loza, Daniel Senif, Scott C. Anderson, David B. Camarillo

## Abstract

Previous research has sought to quantify head impact exposure using wearable kinematic sensors. However, many sensors suffer from poor accuracy in estimating impact kinematics and count, motivating the need for additional independent impact exposure quantification for comparison. Here, we equipped seven collegiate American football players with instrumented mouthguards, and video recorded practices and games to compare video-based and sensor-based exposure rates and impact location distributions. Over 50 player-hours, we identified 271 helmet contact periods in video, while the instrumented mouthguard sensor recorded 2,032 discrete head impacts. Matching video and mouthguard real-time stamps yielded 193 video-identified helmet contact periods and 217 sensor-recorded impacts. To compare impact locations, we binned matched impacts into frontal, rear, side, oblique, and top locations based on video observations and sensor kinematics. While both video-based and sensor-based methods found similar location distributions, our best method utilizing integrated linear and angular position only correctly predicted 81 of 217 impacts. Finally, based on the activity timeline from video assessment, we also developed a new exposure metric unique to American football quantifying number of cross-verified sensor impacts per player-play. We found significantly higher exposure during games (0.35, 95% CI: 0.29-0.42) than practices (0.20, 95% CI: 0.17-0.23) (p<0.05). In the traditional impacts per player-hour metric, we observed higher exposure during practices (4.7) than games (3.7) due to increased player activity in practices. Thus, our exposure metric accounts for variability in on-field participation. While both video-based and sensor-based exposure datasets have limitations, they can complement one another to provide more confidence in exposure statistics.

## Introduction

Concussions are a common injury in contact sports, with an estimated 300,000 sports-related concussions in the United States annually [1]. Concussions lead to short-term neurological deficits [2,3], and a history of concussions have been correlated with increased risk of developing long-term neurodegeneration [4]. Thus, to improve player safety and health, it is imperative we understand the mechanism of concussion. One critical step towards understanding concussion mechanisms is to quantify the exposure of players to potentially injurious head impacts. Many studies have found that players may be more susceptible to sustaining a concussion depending on their exposure to head impacts [5,6], and more recent studies have suggested that exposure to repeat subconcussive head impacts over a career could also increase risk of long-term neurodegeneration [7,8].

To quantify head impact exposure to players on the field, previous studies have focused on instrumenting American football players with wearable sensors [5,9–19] (Table 1). These sensors have proven useful in collecting both head impact count and the kinematics of head impacts from a large number of players. In particular, the Head Impact Telemetry System (HITS) [20,21] has been used extensively to collect data on hundreds of thousands of impacts over the past decade, and exposure data collected from these studies have been used to define injury thresholds and develop helmet testing protocols [22].

**Table 1:** Previous exposure studies.

| Previous Study | Team | Seasons | Sensor | Additional Impact Verification | Impact Count | Impact Severity |
| --- | --- | --- | --- | --- | --- | --- |
| Beckwith et. al. [6] | NCAA Div. 1<br>High School Varsity | 2005 - 2010 | HITS | Exclude <10g peak linear acceleration<br>Exclude impacts outside event time<br>Exclude non-head impacts | 14.7 impacts per event<br>for concussed players | 20.7g median, days without concussion<br>22.5g median, days with concussion |
| Brolinson et. al. [10] | NCAA Div. 1 | 2003 - 2004 | HITS | 10g impact threshold | > 7.2 impacts per event* | 15.3g median |
| Broglio et. al. [11] | High School Varsity | 2007 | HITS | 15g impact threshold<br>Questioned athletes severe impacts | 15.9 impacts per event | 24.8g mean during games<br>23.3g mean during practices |
| Crisco et. al. [12] | NCAA Div. 1 | 2007 - 2009 | HITS | Exclude <10g peak linear acceleration<br>Rigid-body modeling<br>Video correlation of impacts >125g | 6.3 impacts per practice<br>14.3 impacts per game | 20.5g median |
| Duma et. al. [14] | NCAA Div. 1 | 2003 | HITS | 10g impact threshold<br>Video correlation of impacts | >15.0 impacts per game*<br>>7.6 impacts per practice* | 32g mean |
| Kawata et. al. [15] | NCAA Div. 1 | 2015 | i1 Mouthguard | 10g impact threshold | 7.0 impacts per event | 30.3g mean |
| Mihalik et. al. [16] | NCAA Div. 1 | 2010 | HITS | Exclude <10g peak linear acceleration<br>Video correlation of impacts** | 14.4 impacts per game | 24.7g mean in anticipated impacts**<br>27.0g mean in unanticipated impacts** |
| Mihalik et. al. 2 [17] | NCAA Div. 1 | 2005 - 2006 | HITS | Exclude <10g peak linear acceleration<br>Exclude impacts outside event time | 57,024 impacts from 72 players over<br>two seasons | 22.7g mean in full contact practice<br>21.1g mean in games |
| Naunheim et. al. [18] | High School Varsity | Not Reported | Linear Accels<br>in Helmet | 10g impact threshold<br>Video correlation of impacts | 40.5 impacts per player-hour<br>during game | 29.2g mean |
| Shnebel et. al. [19] | NCAA Div. 1<br>High School Varsity | 2005 | HITS | 10g impact threshold | >12.9 impacts per event, college*<br>>21.7 impacts per event, high school* | 58.8g 95th percentile, college<br>56.2g 95th percentile, high school |
| Reynolds et. al. [20] | NCAA Div. 1 | 2013 | x2 Skin Patch | 10g impact threshold<br>Proprietary false impact filter<br>Exclude impacts outside event time | 24.2 impacts per game<br>16.8 impacts per full pad practice | 28.2g mean during games<br>28.8g mean during full pad practices |

\* Papers state impacts per event with at most some number of players equipped in an event. Impact count is calculated assuming all players equipped, but this is not necessarily true and is not reported
\*\* Mihalik et. al. only performs video analysis on a subset of impacts collected and does severity statistics on the subset

However, recent studies have shown that HITS, and many other wearable sensor systems, may suffer from poor accuracy in both classifying head impacts (impact count) [23–25], and measuring head impact kinematics (impact severity and location) [21,26–28]. This motivates the need to verify sensor-based measurements through comparison with other independent exposure measurements.

Video assessment represents an option for collecting an independent dataset of head impact exposure. Video assessment has been used extensively to review plays and sports injuries by coaches and medical professionals [29–33]. Several sensor-based exposure papers also use video to confirm sensor recorded impacts [11–13,16,17]. However, current methodologies assume that sensors are capable of detecting all head impact instances and video assessment is dependent on sensor-based recordings. Without independently analyzing video and sensor measurements, bias could be introduced into the exposure analysis. Furthermore, previous studies have only evaluated sensor head impact precision and do not correct for false negatives in impact detection.

In this work, we introduce a video assessment protocol to collect an independent video-based head impact exposure dataset from seven collegiate American football players for a single season. We also equipped players with an instrumented mouthguard [23,27,34,35], which collected impact kinematics. The main goal of this study was thus to compare the exposure rates and impact location distributions determined by independent video-based and sensor-based methodologies. This will help inform future protocols for quantifying exposure data and using complementary analysis of sensor or video data.

In addition, we combine sensor and video information to introduce a new exposure metric for quantifying head impact exposure unique to American football, which measures the number of head impacts sustained over a single player-play. While traditionally, exposure is quantified as the number of impacts per player exposure (defined as a player participating in a practice or game) or per hour of play time, American football is a unique sport where every player is expected to spend varying amounts of time on the field and in play. As an example, statistics on National Football League games report an average of 133.6 plays per game, of which 10.3 involve punting. Thus, specialized players such as punters are on the field, and thus exposed, far less often than players such as linemen. Our exposure metric accounts for variability in on-field participation, and normalizes exposure by the base unit for American football: a single play. This metric is also a prime example for the complementary analysis of video and sensor exposure because head impact kinematic sensors cannot measure non-impact activities and video does not have sufficient resolution to identify discrete head impacts.

## Methods

To collect a video-based head impact exposure dataset, we relied on a tiered video assessment protocol. Impact exposure and kinematics were also independently measured using the instrumented mouthguard.

### Sensor-Based Exposure Data Collection

Data were collected from the Stanford University American football team during the 2015 fall season. Seven players were consented for study participation through the Stanford Internal Review Board IRB #34943. Players represented a variety of positions (three offensive linemen, one running back, one fullback, one wide receiver, and one defensive linebacker).

Players wore instrumented mouthguards [34,35] (Fig 1) for the entire fall season in all practices and games. Before the 2015 fall season, we collected upper dentition impressions from our seven consented players. Electronic boards containing kinematic sensors, flash memory, and processing were embedded between two layers of 3mm ethylene vinyl acetate, which were pressure formed against the dentitions.

**Fig 1.**
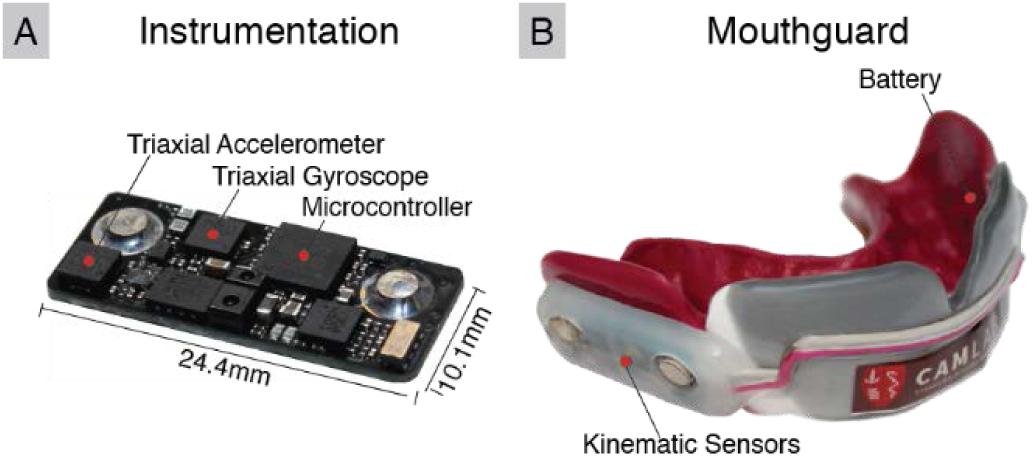
Instrumented mouthguard for measuring impact severity. (A) Sensor board containing tri-axial linear accelerometer, tri-axial angular gyroscope, and infrared proximity sensor are embedded (B) inside a custom-formed instrumented mouthguard.

The electronic boards contained a tri-axial linear accelerometer (H3LIS331) and tri-axial gyroscope (ITG3701A), both sampling at 1000Hz. The linear acceleration data were filtered at CFC180 (300Hz fourth order Butterworth low pass filter) [36] while the angular velocity data were filtered with a 184Hz fourth order Butterworth low pass filter (manufacturer defined bandwidth). Impacts were identified using a 10g linear acceleration magnitude threshold, and data were collected 10ms before and 90ms after impact trigger. Furthermore, the instrumented mouthguard was equipped with an infrared (IR) proximity sensor (AMS TMD2771) designed to detect the presence of teeth within the mouthguard tray [23,37]. This allowed us to only consider impact events where the instrumented mouthguard was properly worn, as players commonly removed the instrumented mouthguard when on the sideline. The mouthguard was designed to last 3.5 hours and collect up to 1638 impacts. While the primary purpose of this work was to collect an independent video-based exposure dataset for comparison against sensor-based exposure statistics, the dataset also serves as a training dataset to design an impact detection algorithm for the instrumented mouthguard [37].

Impacts collected on the instrumented mouthguards were time stamped based on an internal real-time clock to within one second. Internal clocks were synchronized to the National Institute of Standards and Technology time (nist.time.gov) prior to an event. A unique “time-sync” mouthguard not built for any players was used to synchronize mouthguard times with the camera times for impact correlation. This was accomplished by having an athletic trainer apply known impacts on the “time-sync” mouthguard in view of all cameras. The real-time stamps on the “time-sync” mouthguard were matched with those on the cameras after the event.

### Video-Based Exposure Data Collection

Our video assessment protocol utilized multi-angle video to better view players on the field, and a tiered assessment method for both sensitive and specific identification of activities, most notably helmet contact periods (Fig 2).

**Fig 2.**
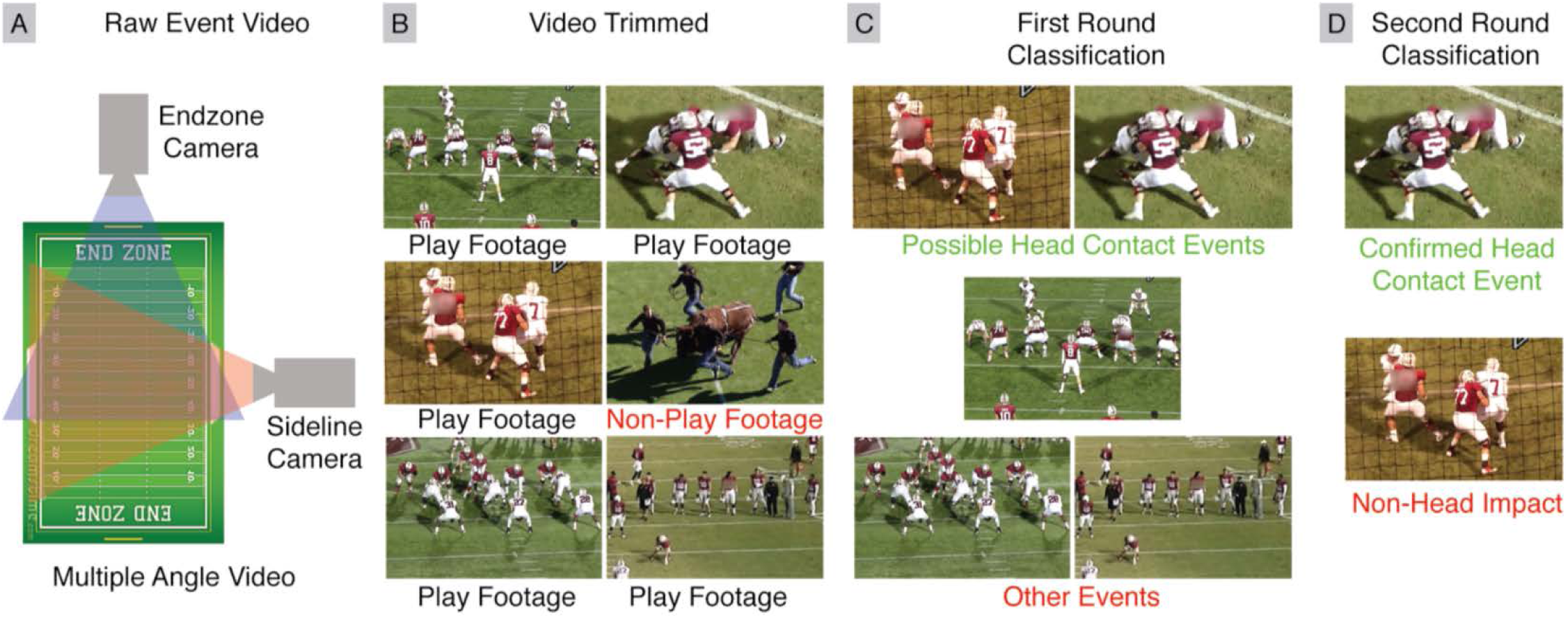
Overview of tiered video assessment for collecting head impact exposure dataset. (A) Multiple video angles were collected for each practice and game, with at least one camera capturing an end-zone view and one camera capturing a sideline view. (B) Video was trimmed by technicians to only include play footage. (C) Trained raters performed a first round of video assessment, tracking specific players and labeling their activity. (D) A second round of video assessment performed by one of the authors confirmed Helmet Contact activities.

Video was first collected during practices and games from multiple viewing angles using 1080p resolution cameras recording at 30fps. We obtained two videos per field, with one video taken from the end-zone and another taken from the sideline. Because practices utilized multiple fields, there were four videos per practice, and two videos per game.

Next, videos were trimmed to only include plays, which cut video time in half. Then, in a first round of video assessment, several trained raters were tasked with tracking single players in each video and tagging their activity. In this first round of video assessment review, raters were instructed to be sensitive to Helmet Contact activities. Finally, one of the authors, who had over three years of experience deploying wearable sensors and evaluating video footage, performed a second round of video assessment of all identified periods of Helmet Contact. The author viewed periods of Helmet Contact from multiple angles with the purpose of being specific in identifying Helmet Contact activities.

### Video Assessment First Round

During the first round of video assessment, raters were tasked with tracking single players in all videos associated with an event and labeling their activity. A total of fourteen raters were trained to classify player actions in the videos.

We classified player activity with six distinct labels: Helmet Contact, Body Contact, No Contact, Obstructed View, Idle, and Not in Video (Fig 3). We defined Helmet Contact activity as any continuous period with direct contact to the tracked player’s helmet, and Body Contact as any continuous period with contact elsewhere on the body. While Body Contact could induce head accelerations through whiplash effects, they were not considered head impacts in this analysis. No Contact events were classified whenever players were observed on the field, participating in a play, but not in contact with anyone or anything. Obstructed View labels were used whenever a player’s helmet could not be clearly seen in video due to obstruction, such as when a player was tackled by multiple opponents and could not be seen. Idle events were classified whenever a tracked player was observed in video, but not on the field participating in the play. This was most commonly observed when the player was on the sideline. Finally, Not in Video events occurred when a player was not visible.

**Fig 3.**
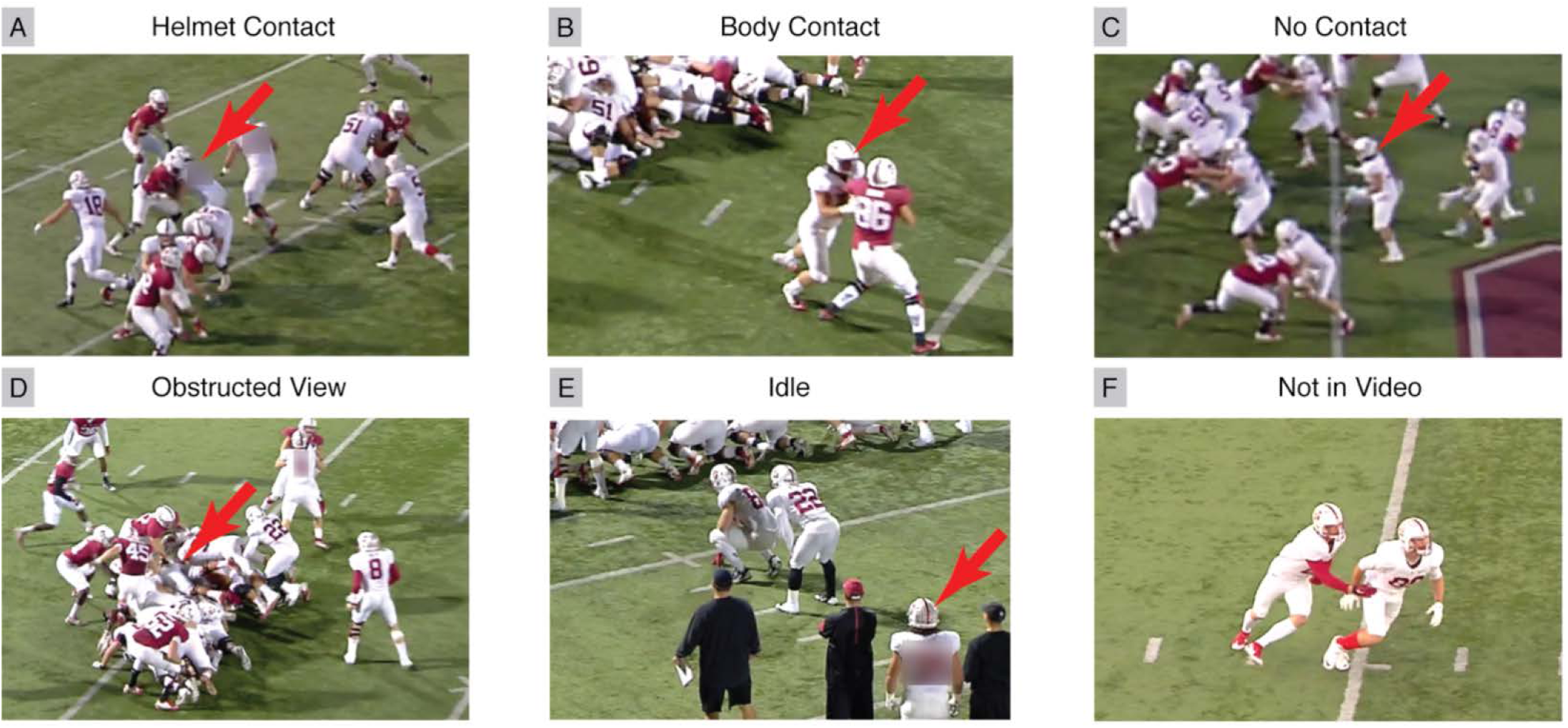
Activity classifications for tracking player activity and identifying Helmet Contact activities with high sensitivity. Tracked player marked with a red arrow. (A) Raters identified Helmet Contact activities whenever the tracked player’s head overlapped with an opposing player. (B) Body Contact activities when there was contact not involving the head. (C) No Contact activities when player was in play, but not actively in contact. (D) Obstructed View activities when there was no clear view of the player’s head. (E) Idle activities when players were observed on the sideline, or otherwise not in play. Finally, (F) Not in Video activities when tracked player was not in the video.

For the purposes of this study, we asked raters to be particularly sensitive to identifying Helmet Contact activities. Helmet Contact activities were tagged whenever the tracked player’s helmet overlapped with another player or object in video. Additionally, multiple Helmet Contact activities were tagged when there were potentially multiple independent helmet impacts in quick succession. This provided an upper bound on the number of Helmet Contact activities sustained.

Due to the large number of videos, each video was only viewed by one rater. Thus, it was necessary to train raters to classify player actions consistently and assess rater reliability. Because we instructed raters to be sensitive to Helmet Contact events, we assessed raters on their reliability in identifying Helmet Contact events. First, raters were trained on a 2-minute demonstration video that explained activity classifications. Next, raters were evaluated on a 10-minute video to ensure good rater reliability. Both training and evaluation video clips were taken from videos collected as part of this study. Rater reliability was assessed by comparing the number of Helmet Contact events classified by each rater in the 10- minute evaluation video to the number of Helmet Contact events classified by the author performing a second round assessment. The fourteen raters classified an average of 88% of the Helmet Contact events identified by the second round assessment author.

### Video Assessment Second Round

After raters classified player actions within the videos, an author with over three years of experience deploying wearable sensors and analyzing video footage performed a more detailed analysis for specific identification of Helmet Contact activities. Because Helmet Contact represents potential head impacts and exposure to injury, it was necessary to perform a more detailed breakdown to confirm a Helmet Contact activity and obtain qualitative information such as contact location and directionality.

For each rater identified Helmet Contact activity, the author reviewed all video angles to confirm Helmet Contact activity (Fig 4). Observed Helmet Contact activities were then further classified by contact location to the player (helmet front, side, top, rear, and front and rear oblique), and contact location on the opponent (helmet, body, upper limb, lower limb, and ground).

**Fig 4.**
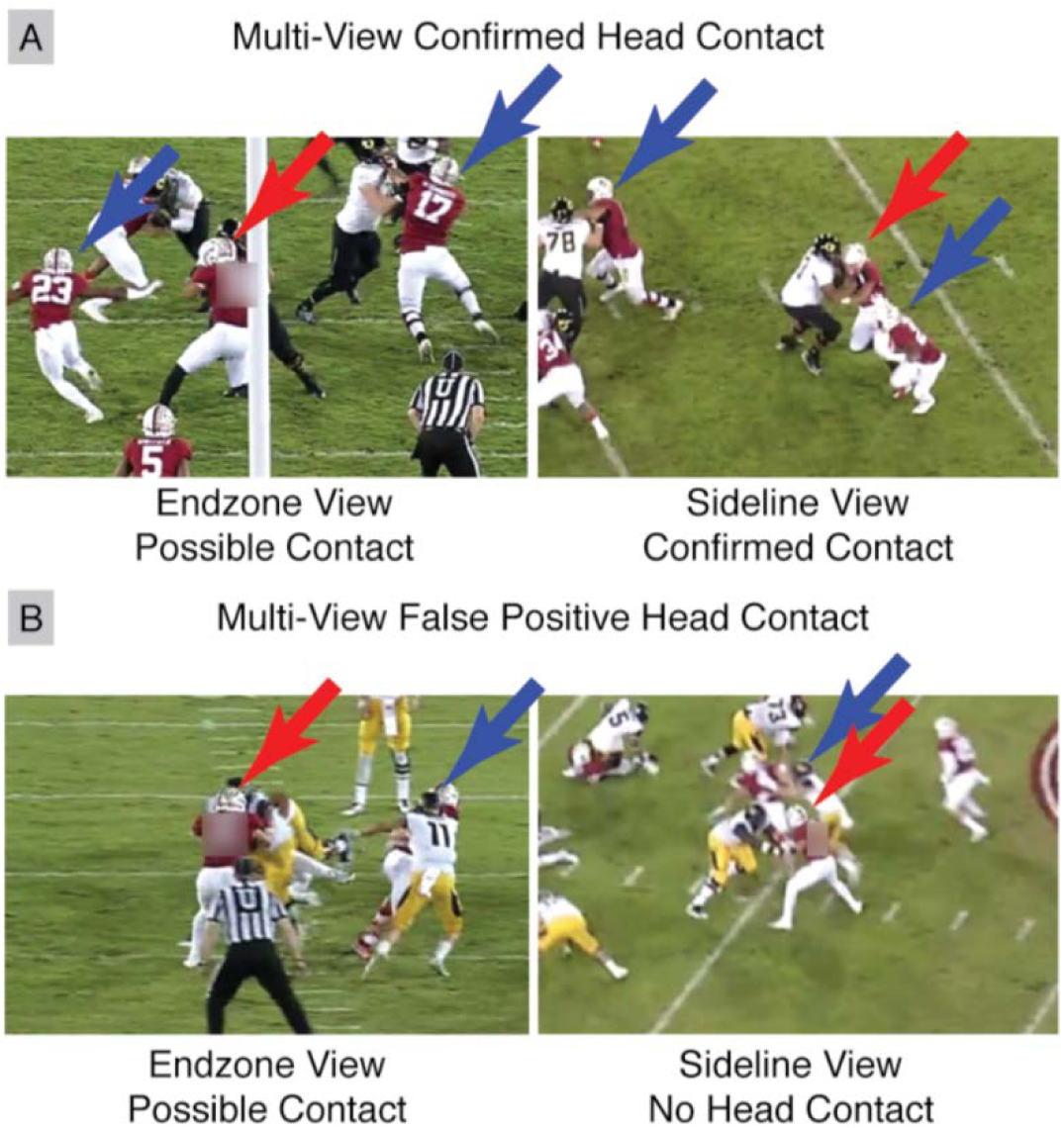
Second round video assessment for specific Helmet Contact activity identification. Multiple videos were used to confirm Helmet Contact activities. Red arrows mark the tracked player, with blue arrows marking other players. End-zone videos show helmet overlap, but sideline video showed (A) definitive head contact and (B) no helmet contact.

### Comparison of Sensor-Based and Video-Based Exposure

Using the sensor and video protocols, we were able to collect independent sensor-based and video-based impact exposure data. Helmet Contact activities were compiled after the second round of video assessment. For the sensor-based head impact exposure data, we took discrete impact recordings from the instrumented mouthguard and considered only head impacts that occurred during practice or game periods (as provided by athletic training staff) and had IR readings indicating instrumented mouthguards were properly worn. We then cross-verified helmet contact periods and discrete impact events between video-based and sensor-based exposure sets respectively by comparing the synced time-stamps. This differs from previous video confirmation protocols in that we identified helmet contact periods in video independently [24,25].

### Comparing Exposure Rates

Using the independent video-based helmet contact periods, sensor-based impacts, and cross-verified impacts, we computed a traditional exposure metric: the number of helmet contact periods or head impacts per player-hour (normalizing by number of hours recorded by both the mouthguard and video). We then computed our new exposure metric: the number of cross-verified sensor-based head impacts per player-play. Plays were discretized using the video-based player activity timeline combined from all videos during an event. Helmet Contact, Body Contact, and No Contact activities were associated with players in play, and the Idle and Not in Video activities were associated with players not in play. Sequences of Helmet Contact, Body Contact, and No Contact activities were represented by a single play. Finally, we also computed the number of plays per player-hour.

### Comparing Impact Location Distributions

Next, we compared the impact location distributions from cross-verified video-based helmet contact periods and sensor-based head impacts. We defined location vectors representing front (1, 0, 0), rear (−1, 0, 0), side (0, ±1, 0), oblique (±1, ±1, 0), and top (0, 0, 1) impacts (Fig 5). During second round video assessment, the author qualitatively classified cross-verified helmet contact periods into a location bin to obtain a video-based impact location distribution. For sensor-based impact location distribution, we explored several processing techniques of the cross-verified head impact kinematics to bin impact locations.

**Fig 5.**
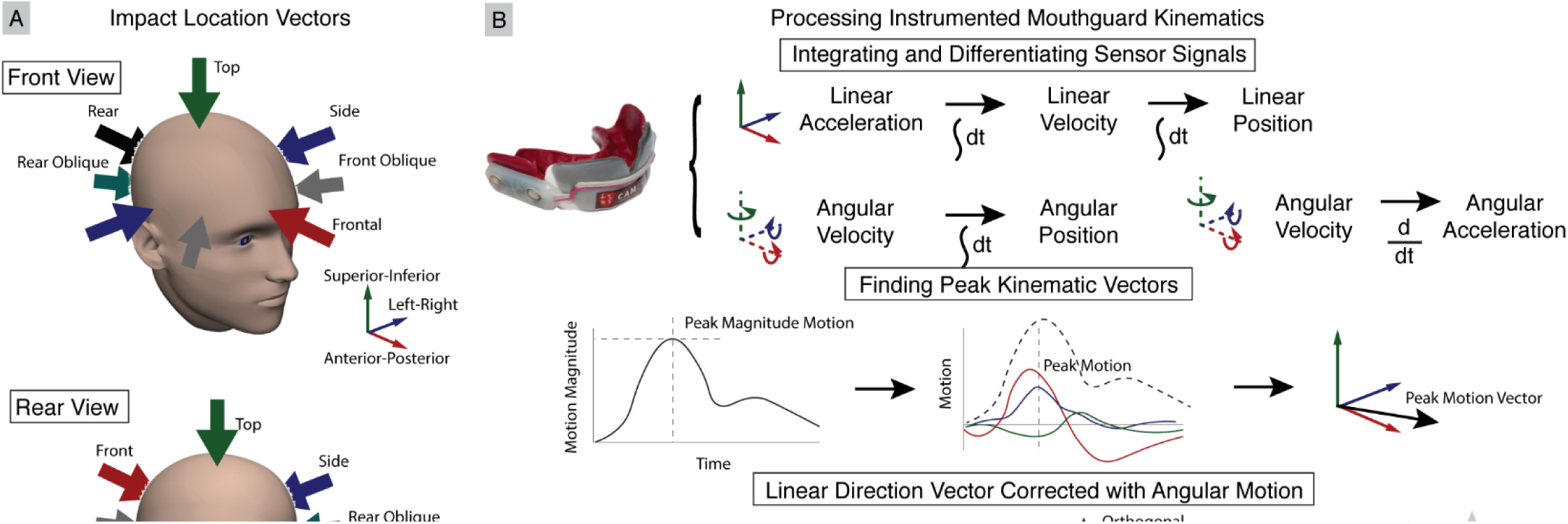
Impact Location Vectors and Mouthguard Kinematics Processing. (A) Locations are binned into front, front oblique, side, rear oblique, rear, and top impacts. (B) Video-based helmet contact periods were qualitatively binned into impact locations during second round video assessment by the rating author. For sensor-based head impacts, kinematics were processed by first integrating or differentiating sensor linear acceleration and angular velocity signals to obtain linear velocity, linear position, angular acceleration, and angular position (represented with XYZ Euler angles). Peak motion (angular or linear acceleration, velocity, or position) vectors were found by identifying the peak magnitude and determining the 3 degree-of-freedom components. Peak linear acceleration, velocity, and position vectors were binned directly. We also incorporated peak angular motion vectors to correct respective peak linear motion vectors.

First, we integrated sensor linear acceleration and angular velocity data to obtain linear velocity, linear position, and angular position (defined using XYZ Euler angles) during each impact. We also differentiated angular velocity data to obtain angular acceleration [27,38]. We then found the time of peak magnitude linear and angular accelerations, velocities, and positions during each impact. Using the time of peak magnitude, we identified the 3 degree-of-freedom peak linear and angular acceleration, velocity, and position vectors. Traditionally, the peak linear acceleration vector has been used to define impact location, with impact location occurring in the opposite direction (an impact to the front of the head causes acceleration towards the rear of the head) [13]. Thus, we binned impacts using linear data by separately finding the location vector with the smallest angle with respect to the sensor-based peak linear acceleration, velocity, and position vectors.

To utilize the peak angular data, we note that if we treat the head and neck as a simple pendulum, the linear and angular motion vectors should be orthogonal. Thus, we define a corrected motion vector by first taking the cross product of the peak linear acceleration, velocity, or position vector with the respective angular acceleration, velocity, or position vector. We then cross the result with the respective angular acceleration, velocity, or position vector to obtain a corrected motion vector, whose location we bin by again finding the location vector with the smallest angle.

The cross-verified discrete impacts from the instrumented mouthguard were also used to assess impact severity exposure to compare against previously reported values. We defined severity based on the peak linear acceleration, angular velocity, and angular acceleration magnitude from each impact.

## Results

Fourteen raters reviewed over 163 hours of video for the seven players spanning 11 practices and 6 games during the 2015 fall season. This represented over 62 hours of mouthguard recordings with over 50 hours of actual in-game or in-practice time. In that time, players participated in 927 plays. Raters identified 1004 Helmet Contact activities after first round video assessment, of which 271 were confirmed following second round video assessment. The instrumented mouthguard recorded a total of 13,034 impacts, of which 10,949 occurred during practice or game periods. 2,032 of these were determined to be in-mouth impacts based on IR readings. When matching time stamps of video-based helmet contact periods and sensor-based head impact events, we cross-verified 193 video-based helmet contact periods and 217 sensor-based impact events (Fig 6).

**Fig 6.**
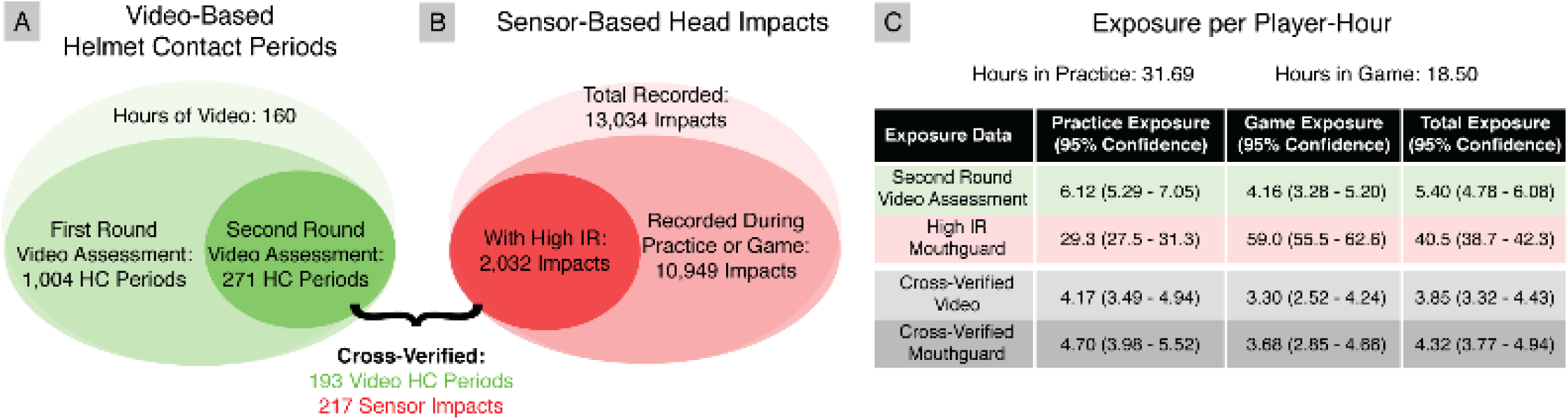
Comparison of Video-Based and Sensor-Based Head Impact Exposure. (A) Exposure rates collected from independent (A) video-based and (B) sensor-based methods differed drastically. Our instrumented mouthguard identified an order of magnitude more discrete head impacts than video-based helmet contact periods. Cross-verifying head impacts with helmet contact periods yields a more consistent 217 discrete head impacts within 193 helmet contact periods. Delineating by event type, we found that there was greater head impact exposure in practices than in games.

### Impact Exposure

Treating sensor-based and video-based exposure data separately, we observe a large discrepancy in exposure rates (Fig 6). The instrumented mouthguard identified 2,032 impacts in the 50 hours of recorded play-time, which gives an exposure rate of 40.5 (95% CI: 38.7-42.3) impacts per player-hour. In video assessment, there were 271 video-based helmet contact periods, which gives an exposure rate of 5.40 (95% CI: 4.78-6.08) helmet contact periods per player-hour. Finally, the 217 sensor-based discrete head impacts cross-verified with video helmet contact periods gives an exposure rate of 4.32 (95% CI: 3.77-4.94) impacts per player-hour.

Next, we further delineate exposure based on event type (games or practices). When considering sensor-based results independently, the instrumented mouthguard had double the head impact rate in games than in practices. For independent video-based exposure and cross-verified exposure, exposure during practices was higher. Even for cross-verified mouthguard impacts, we found greater head impacts per player-hour exposure in practices (4.70, 95% CI: 3.98-5.52) than in games (3.68, 95% CI: 2.85-4.66), though the difference is not significant (p=0.105). When considering the number of cross-verified sensor-based impacts per player-play however, we found that most players sustained more impacts per player-play during games than during practices. Indeed overall, players experienced significantly fewer impacts per player-play during practices (0.20, 95% CI: 0.17-0.23) than during games (0.35, 95% CI: 0.29-0.42) (p<0.05). This discrepancy is likely due to a difference in amount of play seen, where all players participated in significantly more player-plays per player-hour in practices (23.16, 95% CI: 21.52-24.90) than in games (10.43, 95% CI: 9.01-12.01) (p<0.05).

### Impact Location

We quantified the distribution of impact locations in our cross-verified video-based helmet contact periods and sensor-based head impacts (Fig 7). In both cases, the majority of helmet contact periods or head impacts occurred to the front, front oblique, and sides of the head respectively. 40.5%, 23.1%, and 30.3% of impacts were observed in video assessment in the front, front oblique, and sides of the head for a total of 93.8% of impacts. According to our processed mouthguard kinematics, an average of 85.8% of impacts occurred to the front, front oblique, and sides of the head.

**Fig 7.**
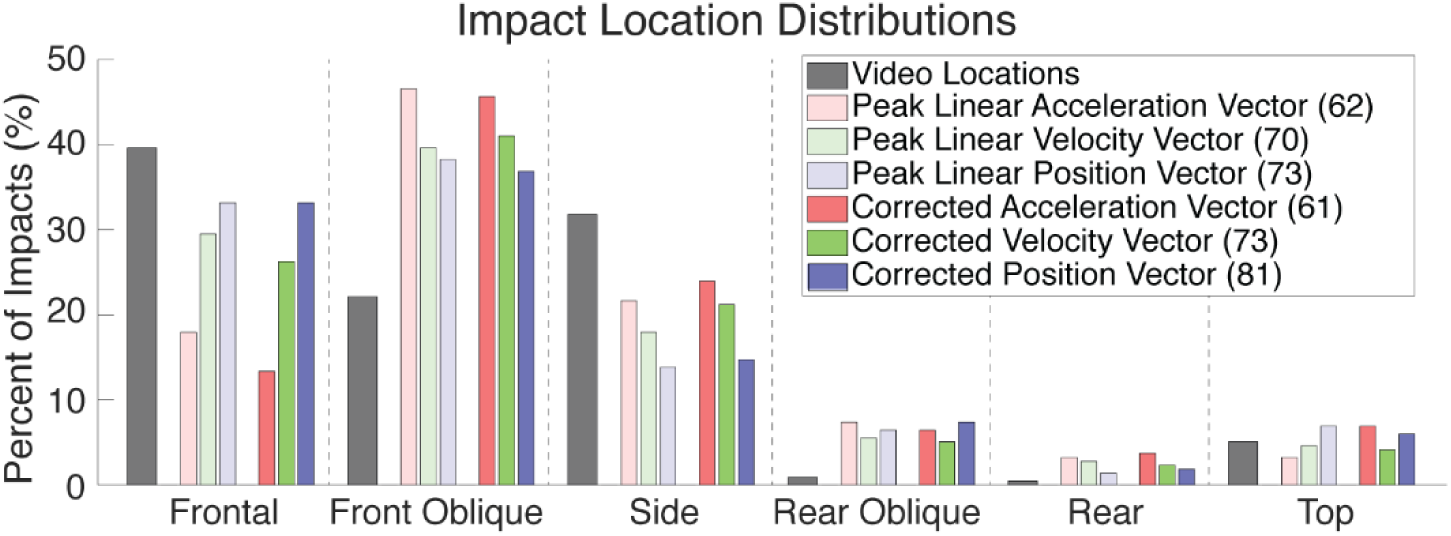
Comparison of Video-Based and Sensor-Based Impact Location Distributions. For both video-based and sensor-based distributions, the majority of impacts are to the front, front oblique, and sides. Methods for processing sensor kinematics to obtain location generally did not match well with video locations (number of matches in parentheses). Methods using integrated (position) kinematics and incorporating angular motions (corrected) had the best match.

When comparing the location determined through video with the location determined used the processed mouthguard kinematics, we found that our best processing technique (corrected linear position) only predicted impact location correctly in 81 of 217 impacts (37.3%). The traditional peak linear acceleration process, however, was least accurate with only 61 of 217 impacts (28.1%) correctly predicted. In general, correctly identifying impact location improved with integration, and improved when assuming orthogonality with angular measures.

We also quantified head impacts based on the opposing impact object (opposing helmet, body, limb, or ground), which can only be determined from video assessment. We found 64% were to an opponent’s helmet, 27% were to an opponent’s body, 5% were to an opponent’s upper or lower limbs, 3% were to the ground, and 1% were to other objects such as practice pads.

### Impact Severity

Impact severity was only quantified in the 217 impacts recorded by the instrumented mouthguard with high IR and cross-verified with video. We delineate impact severity based on event (games or practices). Over all players, the median peak linear acceleration, angular velocity, and angular acceleration magnitude was 18.4g (95% CI: 15.2g-19.8g), 9.4rad/s (95% CI: 8.5rad/s-10.2rad/s), and 1240.5rad/s^2^ (95% CI: 1109.1rad/s^2^-1475.9rad/s^2^) respectively during practices, and 23.2g (95% CI: 18.5g-26.1g), 12.5rad/s (95% CI: 10.8rad/s-13.9rad/s), and 1435.0rad/s^2^ (95% CI: 1288.4rad/s^2^-1723.3rad/s^2^) respectively during games. While the medians were larger during games, only peak angular velocity magnitude had a significant difference (p<0.05).

## Discussion

In this work, we present a tiered video assessment protocol for obtaining an independent video-based dataset for head impact exposure in American football to compare against sensor-based impact exposure and impact location distributions. This work differs from previous head impact exposure datasets that rely primarily on imperfect head impact sensors, with a few datasets further confirming true positive vs. false positive head impacts in video. Instead, we treat the video assessment as an independent exposure dataset. Exposure datasets collected with impact sensors alone are limited by a high occurrence of false positives due to the sensitivity of motion sensors, and known innaccuracies of many sensor systems. Video assessment of exposure data are limited by the qualitative nature of analyzing video and the inability to quantify impact kinematics. However, in this study, we demonstrate that the two methods can complement one another and reveal directions for improvement in sensor-based and video-based data collection. Combining these information, we also introduce a new American football specific exposure metric based on the number of sensor-based discrete head impacts per video-observed player-play, which account for players varied involvement during events.

### Comparing Independent Exposure Data

To demonstrate the limitations of independent video-based and sensor-based head impact exposure, we highlight key differences between between the independent exposure sets and the exposure set determined with impacts cross-verified in both video and sensors.

First, we note that the instrumented mouthguard only detected 71.2% of the video classified Helmet Contact events, missing 76 of 271 events. One possible reason for this is that the instrumented mouthguard was set to record an impact event when the linear acceleration magnitude exceeded 10g, which is similar to impact thresholds employed in previous work (Table 1). In video assessment however, Helmet Contact events were identified by observing physical contact of the helmet with another object, not accounting for the severity of the contact. Thus, observed Helmet Contact events could have linear accelerations below the 10g sensor threshold. In fact, 81% of unrecorded impacts were noted as “light severity” or involved body or facemask impacts.

Previously, the 10g sensor threshold was chosen to differentiate head impacts from other events resulting in head motion, such as jumping or running [39]. But, as researchers are suggesting multiple mild impacts could have implications on brain injury risk and long-term degeneration, it is becoming increasingly important to identify any head impact, regardless of severity. Moving to a 20g threshold would result in a loss of an additional 103 head impacts from our dataset, resulting in a sensitivity of 34.0%. This agrees with previous literature investigating the importance of the linear acceleration threshold in identifying impacts [39].

Second, the instrumented mouthguard alone detected 2,032 impacts, which was an order of magnitude greater than the number of cross-verified impacts. This demonstrates the relatively high rate of false positives that have plagued similar impact sensors in the past, and are commonly mitigated using video confirmation (Table 1). On the other hand, we note that it is difficult to observe discrete impact events in video, and that video can only identify contact periods during which multiple discrete impact events could occur. In our cross-verified dataset, we matched 217 discrete sensor-based impacts with 193 video-based helmet contact periods, meaning that up to 13% of video-based helmet contact periods contained multiple discrete impact events.

In addition to differences in exposure, we also discuss discrepancies in impact location distributions between video-based and sensor-based data. In video, we found that 93.8% of impacts occurred to the front, front oblique, and side of the head after the second round of video assessment. Processing instrumented mouthguard kinematics found similar trends for impact location with an average of 85.8% of impacts occurring in the front, front oblique, and side locations; however, the mouthguard impact location matched video identified location in only 37.3% of the impacts using the best corrected position process. Current methods for predicting impact location from impact kinematics are relatively primitive, with peak linear acceleration being the most common method [13]. Peak linear acceleration was the worst at predicting video-based impact location in our dataset. Instead, integrated measures (linear velocity and position) performed better as they account for holistic motion of the head over the entire impact period. Furthermore, incorporating angular measures also improved prediction of video-based impact location, as angular motion provided another source of kinematic information consistent with expected head and neck biomechanics.

Despite some improvements using integrated signals and angular kinematics, matching between video-based and sensor-based impact location was still relatively poor. This could be the result of decoupling between helmet impact location observed in video, and head motion as measured by the mouthguard. In a direct impact, the impact location is expected to correspond with the direction of motion, however we note that in some cases, our raters observed impacts in which helmets “slid past one another”, resulting in impact locations that do not necessarily correspond to the measured direction of motion. Furthermore, identifying impact locations in video is subjective in nature. When relaxing the criteria for matching video-based and sensor-based impact locations to allow for neighboring impact locations (e.g. frontal and frontal oblique), all sensor processing methods were able to predict similar video-based impact locations in 140-150 of 271 impacts.

In addition to discrepancies in head impact location between video-based and sensor-based datasets, our observed distribution of head impact locations differs with previously reported impact location distributions, which report up to 31.9% of impacts to the rear of the head [40]. The discrepancy in impact location distributions may be a result of inaccuracies of previous head impact sensors. From Table 1, we see that the majority of previous exposure studies rely on HITS, which has been shown in the laboratory to have kinematic inaccuracies. Of note, HITS has also been shown to misclassify impacts to the facemask as rear impacts, which may skew impact location distributions [41].

### Head Impacts Exposure per Play

To take advantage of both our sensor-based and video-based exposure data, we present a novel head impact exposure metric specific to American football. For our exposure metric, we look specifically at the number of sensor-based head impacts normalized by the number of video-based player-plays, to account for variability in on field activity between different player positions. Schnebel et. al. alluded to head impact exposure per player-play, stating that linemen tend to have head impacts every play whereas skill players have head impacts more rarely [18]. We have utilized this new metric because of the unique way in which American football is played, where players are not on the field for the entire game.

Table 2 demonstrates the power of reporting head impact exposure using the metric. First, we compare the head impact exposure of a linebacker in two separate games, in which he was observed for 90-120 minutes. In the first game, the linebacker had only 3 head impacts in 6 plays, whereas in the second game, the linebacker had 14 head impacts in 35 plays. In this example, we see that the linebacker had a near 4-fold increase in head impacts per player-hour. However, the per play exposure was more consistent across the two games.

**Table 2:** Head impacts per play exposure metric.

| Player ID | Game Date | Time Observed | # of Plays | # of Impacts | Impacts per Player-Hour | Impacts per Player-Play |
| --- | --- | --- | --- | --- | --- | --- |
| Same Player, Different Games |  |  |  |  |  |  |
| 411 Linebacker | 11/07/2015 | 1:28:52 | 6 | 3 | 2.0 | 0.50 |
| 411 Linebacker | 11/21/2015 | 1:50:39 | 35 | 14 | 7.6 | 0.40 |
| Different Players, Same Game |  |  |  |  |  |  |
| 237 Lineman | 11/28/2015 | 1:14:00 | 30 | 20 | 16.2 | 0.67 |
| 751 Lineman | 11/28/2015 | 1:34:25 | 5 | 3 | 1.9 | 0.60 |

In the second example, we observe two different linemen participating in the same game. The first lineman is a starting lineman, whereas the second lineman is only used in special packages. We see that while both players were recorded for 75-95 minutes, the starting lineman sustained many more head impacts. In terms of head impacts per player-hour, the starting lineman had a more than eight-fold increase compared with the second lineman. Again, the per play exposure is more consistent across the two linemen. We believe this metric of defining head impact exposure in American football prevents skewing from variable on-field participation. We note that while plays are well-defined during games, they are not as clear during practices. Delineating plays in the video activity timeline was done automatically based on video assessment datasheets and may over-estimate the number of plays in practice due to the use of fast and short individual drills which were commonly counted as a single play. The metric can be further refined to properly count practice drills, though we note that practice drill time is currently counted in per-hour or per-event exposure statistics. Similar metrics accounting for player participation may also be useful for sports like ice hockey and basketball where players also spend varying amounts of time in play.

With this exposure metric, we found that there were significantly more head impacts per player-play during games (0.35, 95% CI: 0.29-0.42) than during practices (0.20, 95% CI: 0.17-0.23). From instrumented mouthguard kinematic measurements, we found median head impact linear acceleration, angular velocity, and angular acceleration magnitude was 18.4g (95% CI: 15.2g-19.8g), 9.4rad/s (95% CI: 8.5rad/s-10.2rad/s), and 1240.5rad/s^2^ (95% CI: 1109.1rad/s^2^-1475.9rad/s^2^) respectively during practices, and 23.2g (95% CI: 18.5-26.1g), 12.5rad/s (95% CI: 10.8rad/s-13.9rad/s), and 1435.0rad/s^2^ (95% CI: 1288.4rad/s^2^-1723.3rad/s^2^) respectively during games. Together, these results suggest that head impacts are both more frequent and more severe in games than in practices, which agrees with previous exposure data.

### Limitations

While we successfully collected an independent video-based exposure dataset utilizing a tiered video assessment protocol, this study has several limitations. First, video assessment takes a substantial amount of time. Analyzing 160 hours of video required 1000 man-hours, primarily due to the rigor and detail required. This prevents manual video assessment from being viable for everyday use. However, this protocol can be used to collect datasets with which to improve sensor processing algorithms for head impact detection and location identification, and automated video assessment based on these protocols can be developed for a more streamlined post-event analysis [42,43]. In fact, the dataset collected here is being used to develop an impact detection algorithm for the instrumented mouthguard [23].

Second, video quality (1080p at 30fps) and quantity (two views per field) also affected the subjectivity of video classification. To account for uncertainty due to poor video quality, we had instructed raters to be sensitive to any potential instance of Helmet Contact, and labeled any helmet overlap with another object as a Contact event. Multiple views were only considered in second round of video review, which accounted for the substantial loss of Helmet Contact events between the first and second round of video assessment (1004 Helmet Contact events following first round, 271 Helmet Contact events following second round). Improved video quality and quantity would increase certainty in video labels. While some institutions can afford superior video quality and quantity, we believe our equipment is a reasonable representation of what many collegiate and lower-level institutions currently use.

Third, we have shown previously that the instrumented mouthguard design can be susceptible to lower mandible disturbances, resulting in overestimation of the measured kinematics [35]. Despite this, impact severity trends agree with previously reported values. Future studies will use an updated instrumented mouthguard design that mitigates these disturbances to obtain more accurate impact kinematics. The impact count data reported in this study does not rely on instrumented mouthguard kinematics, and thus are not affected by measurement errors.

Fourth, we note that our sample population of seven players from a single team during a subset of practices and games in a single season is relatively small. Thus, the purpose of this study was not to generalize exposure data but describe differences in video-based and sensor-based datasets. Despite the small sample size, exposure rates and kinematic magnitudes fall within the range of previously reported American football exposure. Because of the small sample population, we did not include delineation of exposure by player position in the main manuscript, but have included it as supporting information (Figs S1, S2, and S3) with the caveat that most positions were represented by a single player.

Finally, we do not consider Obstructed View activities as head impacts. In some activities labeled as Obstructed View, the tracked player is taking part in a large tackle on the field and is hidden beneath other players. This is particularly true of linemen. It is possible that we miss head impacts during these plays, and it is possible that these plays produce more impacts in under-represented locations (rear, rear oblique, and top). In our analysis, there were 29 observed instances of an Obstructed View.

### Implications for Head Injury Research

In this paper, we have demonstrated through a small field study, the comparison and combination of independent sensor-based and video-based measurements of head impact exposure. Currently in head injury research, many sensor-based approaches are being developed for impact exposure studies. While many sensors have demonstrated promise in enabling on-field measurements of head kinematics, some remaining limitations in kinematic accuracy and impact detection accuracy lower the confidence in estimating impact exposure using a single sensor. As such, many researchers have begun using additional information, such as video verification, to improve the quality of sensor-based measurements, and a working group in the National Institutes of Health are recommending video verification as a component in field deployments [44]. As shown in the current study, video-based measurements or sensor-based measurements alone each have their own limitations, but combining and comparing information from these independent measurements helps to overcome these limitations and increase confidence on the exposure dataset. While video-based analyses are currently too time-consuming and require substantial manpower to be a practical long-term solution in head injury research, high quality exposure datasets generated using combined information from multiple independent sources can serve as training datasets to improve sensor algorithms in their accuracy to estimate exposure. Furthermore, as an extension of the current study, adding additional independent sensor measurements (e.g. having multiple sensors at different locations on the head) in field studies may further help increase confidence on the kinematics measurements of the sensor. As such, it is our suggestion that future field studies of head impact exposure should 1) employ multiple sources of measurements to ensure data quality, especially when individual sensors have not been fully validated or have limited accuracy, and 2) utilize high quality, high confidence exposure datasets to help further validate head impact sensors as a longer term solution in impact exposure measurement.

## Conclusions

In conclusion, we have developed a tiered video assessment protocol for collecting an independent impact exposure dataset in American football. This video assessment protocol can be implemented in other contact sports. We have also developed a new head impact exposure metric for American football based on the number of head impacts occurring per player-play. We have found players sustain significantly more impacts per player-play in practices (0.20) than in games (0.35) (p<0.05). This method can provide additional independent exposure information to verify exposure data gathered using wearable sensors that are not fully validated. The verified exposure information can be used to better manage players so that they are not over-exposed to potentially dangerous head impact events.

### Research Involving Human Participants

All procedures performed in studies involving human participants were in accordance with the ethical standards of the institutional research committee and with the 1964 Helsinki declaration and its later amendments or comparable ethical standards. This study was approved by the Stanford University Internal Review Board IRB #34943. Informed consent was obtained from all individual participants included in the study.

## Acknowledgements

We would also like to acknowledge the Stanford football team, athletic training, and sports medicine with their assistance in this study. In addition, we would like to acknowledge our video raters: Ashwin Bhumbla, Cortlandt Bursey-Reece, Helen Gordon, Rosa Hamaleinan, Manpreet Kuar, Edgar Loza, Michelle Mann, Patrick McColl, Abbie McNulty, Jodie Sheffels, and Wilson Tam.

