## Supplementary Materials for "Comparison of video-based and sensor-based head impact exposure"

**Supporting Information**

For supplemental information, we provide impact exposure, location distribution, and severity delineated by player position. We make the caveat that all positions except for linemen were represented by a single player, and some positions had relatively few impacts with which to perform analysis.

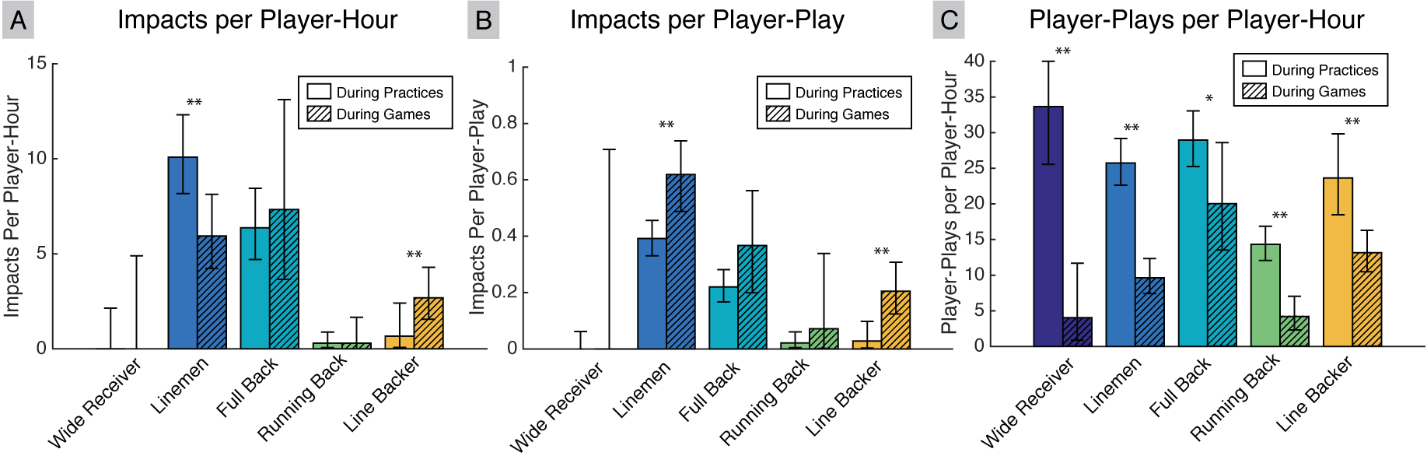

**Fig S1. Impact Exposure by Player Position.** First, we compare exposure across player positions. We compared the exposure data defined by (A) cross-verified sensor-based head impacts per hour and (B) cross-verified sensor-based head impacts per play. We use cross-verified sensor-based head impacts as we believe sensor-based head impacts are the best representation of discrete head impacts and cross-verification gives us high confidence for these impacts. We note that linemen experienced higher exposure than other positions. We also note that, as with the aggregated statistics presented in the main manuscript, players generally experienced more impacts per hour in practices, but more impacts per play in games. (C) This is likely because players participated in more plays per hour during practices than games (* p < 0.05, ** p < 0.01).

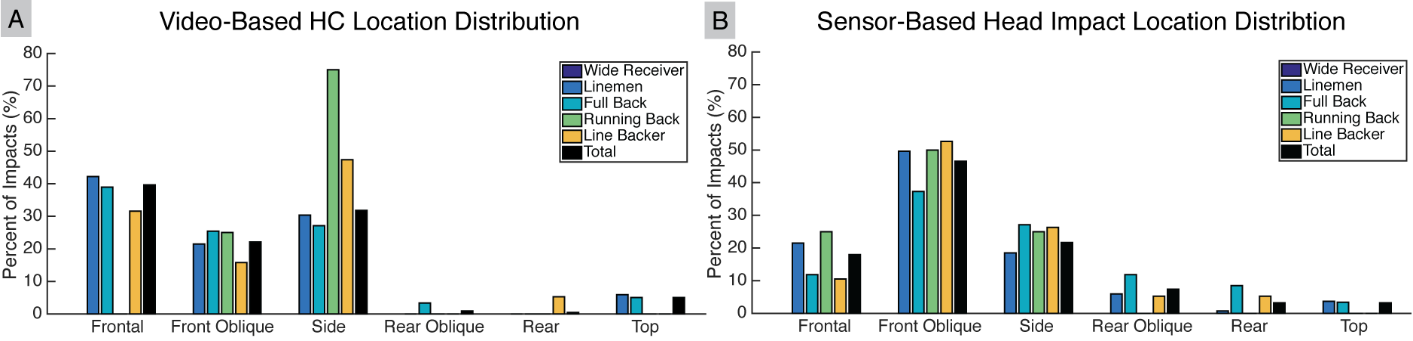

**Fig S2. Impact Location Distribution by Player Position.** We delineated impact location distributions by player position for both cross-verified (A) video-based helmet contact periods and (B) sensor-based head impacts. We used the most common peak linear acceleration vector method for sensor-based head impact location distribution for comparison to previously reported distributions. Players generally have similar impact location distributions in both datasets, though we note that all positions except the linemen are represented by a single player, and some positions observed relatively low exposure.

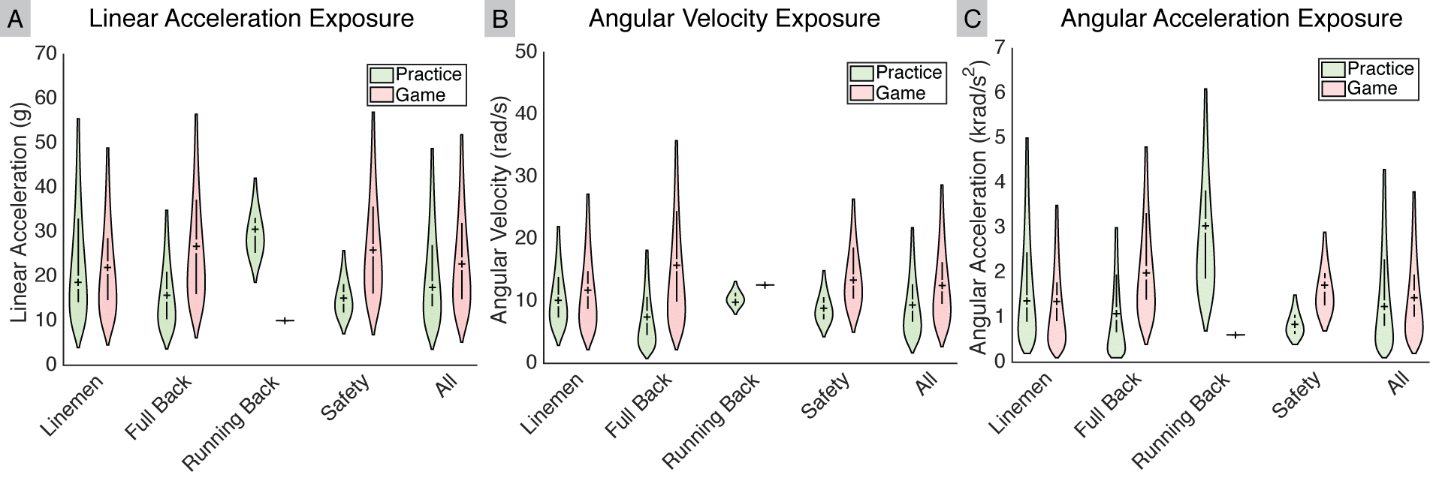

**Fig S3. Impact Severity by Player Position.** Finally, we delineated impact severity by player position. Impact severity were determined from cross-verified sensor-based impacts and were defined with the impact kinematics: (A) peak linear acceleration magnitude, (B) peak angular velocity magnitude, and (C) peak angular acceleration magnitude. The violin plots represent the kinematics distribution observed and were fit using a log-normal distribution. In general, impact severities were higher during games than during practices for all players. Furthermore, while linemen experienced higher exposure rates, they experienced relatively low impact severities.
